# Lower engagement of cognitive control, attention, modulation networks and lower creativity in children while using ChatGPT: an fMRI study

**DOI:** 10.1101/2025.11.07.687207

**Authors:** Tzipi Horowitz-Kraus, Yoed N. Kenett, Dafna Link, Emeel Ashqar, Rola Farah

**Author notes:** Corresponding author: Tzipi Horowitz-Kraus, PhD, Educational Neuroimaging Group, Faculty of Education in Science and Technology, Technion – Israel Institute of Technology, Haifa, Israel; Faculty of Biomedical Engineering, Technion – Israel Institute of Technology, Haifa, Israel.

## Abstract

Generative AI can scaffold idea generation, but how it engages control, attention, and memory systems in the developing versus mature brain, and how this relates to creativity, remains unclear. We compared 6-7-year-old (N = 15) children and adults (N = 16) during a naturalistic 5-minute co-creative dialogue with ChatGPT using the same prompt, in-scanner conversation, undergoing functional MRI scanning and their dialogues were recorded. Functional connectivity (Fisher’s z) was computed within and between seven large-scale networks: cingulo-opercular, default mode (DMN), memory retrieval, frontoparietal (FP), salience, ventral attention (VAN), and dorsal attention (DAN). Creativity scores, assessed outside of the scanner, did not differ between groups. Adults showed stronger within-network connectivity in control/attention systems, most robustly in FP, with additional effects in salience and DAN. Adults also exhibited greater cross-network integration, particularly FP-salience and FP-DAN couplings, alongside broader adult-greater effects among memory- and attention-related pairs. During a co-creative AI-assisted conversation, children exhibit a less coherent and integrated control–attention–memory architecture than adults. The combination of similar creativity scores with divergent neural profiles suggests that developmental differences in the implementation of creative cognition may be more pronounced than baseline capacity. Our findings highlight FP-centric pathways and their salience-gated interactions as candidate substrates by which creativity supports goal-directed co-creative dialogue with AI.

**Highlights:**

- Children demonstrate a lower within-network connectivity in the frontoparietal, salience, and dorsal attention systems while using ChatGPT compared to adults.
- Reduced between-network integration in cognitive control and modulation networks (frontoparietal–salience and frontoparietal–dorsal) in children vs adults while using ChatGPT.
- In adults, but not in children, higher creativity was associated with stronger connectivity within cognitive control networks.

## Introduction

Digital tools have become deeply embedded in children’s daily lives worldwide. Research shows that these technologies, ranging from tablets and laptops to smartphones and emerging generative AI, have a multifaceted impact on children’s learning, well-being, and development^1–7^. While digital tools may offer substantial opportunities to enhance educational experiences, they also introduce challenges such as distraction, cognitive overload, social-emotional strain, and negative impact on literacy development ¹.

In many countries, fostering digital competence and inclusion is a strategic priority, with a vision that this implementation will promote a resilient, innovative, knowledge-based society and will spark creativity. Technology can support learning, but it cannot lead it. Digital tools are not a substitute for educators; they must be used in partnership with them. Children learn best when accompanied by someone who guides, encourages, challenges, and believes in them, a feeling often evoked by generative AI tools^8^. However, how much children are actually engaged cognitively and creatively when interacting with Chat-GPT is yet to be known, which is the topic of the current study.

### Generative AI use by children

Children’s interactions with generative AI (GenAI) are emerging as both opportunities for creativity and challenges for critical engagement. Experimental studies have shown that tools like ChatGPT can support young children’s storytelling by providing interactive, adaptive dialogue ^8^. For example, a study of 5–6-year-olds found that children enjoyed storytelling with ChatGPT, which provided more frequent positive feedback and open-ended prompts than parents, suggesting potential for supporting children’s creative expression when parental time is limited ^8^. Similarly, interventions using games such as *Bot VoyAIge* engaged 10–12-year-olds in critically examining ChatGPT’s responses, revealing that children could identify gender, cultural, and age biases embedded in its outputs ^9^. These findings highlight that while children can find GenAI engaging and even preferable in certain contexts, structured support is necessary to help them navigate its biases and limitations.

Beyond creative engagement, researchers emphasize the developmental, societal, and ethical implications of children’s growing exposure to GenAI. Cross-national work applying a child rights framework has underscored the need to prioritize children’s voices and experiences in shaping AI governance, pointing to rights such as privacy, safety, participation, and development that are often overlooked in current policy debates ^10^. Broader reviews caution that AI may reshape learning, attention, and social interactions, particularly during sensitive developmental windows, requiring careful regulation and proactive safeguards ^11^. Educational interventions that build AI literacy, such as teaching middle schoolers to recognize deepfakes and misinformation, show promise in fostering critical awareness of GenAI’s socio-technical dimensions ^12^. Collectively, these studies demonstrate that children can be both active creators and critical evaluators of GenAI, but ensuring safe, equitable, and developmentally appropriate use will require interdisciplinary collaboration and child-centered policy.

Many fields, including the field of creativity research, are now experiencing and contending with the rise of AI models such as ChatGPT ^13^ which are becoming increasingly popular tools in many linguistic Human-Computer Interactions. The emergence of publicly accessible AI models such as ChatGPT has given rise to global conversations on the implications of AI capabilities and the potential for co-creative cooperation between human and machine agents ^14,15^. Co-creativity is generally defined as multiple parties contributing to a creative process in a blended manner ^16^. Co-creativity is becoming especially relevant in the age of generative AI models ^14^, where highly advanced generative AI can now begin to match humans in creativity tests ^15^. However, increasing evidence highlights not only the benefits ^17–20^, but also the potential costs of AI on human creativity ^21–25^. Thus, elucidating potential differences between children and adult co-creative interactions is greatly needed, and especially elucidating the neural mechanisms that relate to such potential differences in co-creative acts. This, as children’s development continues to shape large-scale neural wiring and may be affected by the rapid AI technological immersion.

### Brain development

A recent study utilizing Electroencephalogram (EEG) examined the cognitive engagement in adult individuals who were instructed to write an essay under several conditions: while using ChatGPT, search engines, or without any assistance. Another condition allowed those who used ChatGPT to write without any assistance, and those who wrote without any assistance were allowed to use ChatGPT for writing ^26^. EEG coherence data demonstrated a cognitive underengagement in ChatGPT users: For writing assignments, individuals were found to engage fewer cognitive resources than individuals who were using other tools to gain knowledge (such as Google) ^26^. Adults, however, have well-developed cognitive, attention, and modulation systems ^27–29^. Neurobiologicaly, brain networks supporting these cognitive control abilities include the frontoparietal and congulo-opercular (Executive Functions; EF system), Dorsal and Ventral Attention Networks (DAN, VAN, bottom-up attention networks), and the modulation system (Default Mode Network; DMN, and the Salience network) ^30^,reach their full developmental maturation at the age of 26 years ^31–33^. The long-range connections and the efficiency of communication between these networks increase markedly through development, reaching maturity in adulthood ^34^. While generating new ideas, as happens when individuals communicate or when they speak to generative AI, there is a need to plan, inhibit, and remember what was already discussed (i.e., working memory)-all parts of EF, which are not yet mature among young 6-year-old children^35^. However, creativity is also important for generating new ideas ^36^. What are the neural mechanisms that support the involvement of EF during a co-creative task such as an interaction with ChatGPT, how do they differ in children versus adults, and how might they be impacted with relation to interaction with ChatGPT?

### Creativity in children

Creativity is broadly defined as the ability to generate ideas that are both novel and useful within a given social context ^37–39^. This construct relies on cognitive processes, including divergent thinking (DT), which encompasses fluency, originality, and flexibility ^40,41^, memory ^42,43^, cognitive control ^44^, attention ^45^, and associative thinking ^46^. Developmental perspectives emphasize that creativity is not a fixed trait but an emergent process, with executive functions (EF) providing regulatory mechanisms to support idea generation and sustain attention ^47^.

However, empirical findings suggest that young children’s DT may rely heavily on bottom-up associative processes, with EF sometimes playing an inverse or constraining role ^41^. Critically, current research focusing on the creative process emphasize dull-process theories, mainly those including creative idea generation and idea evaluation stages ^42,48–50^. A growing scientific focus regards creative idea evaluation, and how it is related to metacognitive capacities, which develop later in life ^51^. Longitudinal studies show substantial gains in fluency and originality between ages 4 and 6, supported by both memory retrieval and mental operations, but also highlight a potential “slump” in creativity when children enter first grade and encounter more formalized, convergent instructional practices ^40^.

The assessment of creativity has often relied on DT tasks such as the Alternative Uses Task (AUT), which capture both product-level measures (e.g., fluency and originality) and process-level mechanisms underlying idea generation ^52^. These tasks are advantageous in their sensitivity to developmental change and in revealing not only the quantity but also the quality of children’s ideas. Complementary measures of evaluative ability are also necessary, as they assess whether children can judge the originality or appropriateness of their own ideas ^52^. Such current assessment tools are based on computational methods such as large language models to provide automatic and quantitative neasures of originality and appropriateness of responses in open-ended creativity tasks such as the AUT ^53–56^.

Contemporary accounts highlight that creativity is shaped by social interaction and cultural context, positioning it as a developmental process embedded in children’s everyday environments ^37,57^. While studies using AI with children demonstrate that generative models can scaffold storytelling and imaginative play, the extent to which such tools enhance or constrain DT remains insufficiently understood, and whether cognitive processes are engaged equally as in adults further requires empirical work.

### The neuroscience of creativity

Over the past few decades, extensive neuroscientific research has been conducted to elucidate the neural mechanisms that realize creative thinking ^58,59^. Such research has been conducted via multiple neuroscientific methods ^60^, and extensively via functional MRI research to elucidate cuntional brain connectivity that supports creativity ^61–65^.

Functional magnetic resonance imaging (fMRI) has significantly advanced the understanding of creativity by highlighting how distributed brain networks support creative cognition. Recent studies have emphasized functional interactions among large-scale brain systems composed of multiple regions. This approach has identified several key networks—specifically, the EF system, and the modulation one (DMN, and salience network)-whose coordinated activity underlies various facets of creative thinking ^63^.

The DMN, as part of the modulation system, comprising regions such as the medial prefrontal cortex, posterior cingulate cortex, angular gyrus, and hippocampus— has emerged as central to creative cognition due to its role in memory retrieval and associative thinking ^66^. The DMN supports memory processes essential to creative ideation, especially remote associative thinking—the ability to link conceptually distant ideas ^46,67,68^. Beyond idea generation, the DMN also contributes to evaluating ideas by comparing them to stored knowledge to determine novelty and by mentally simulating their feasibility to assess appropriateness ^42^. The salience network is comprised of the anterior insula and dorsal anterior cingulate cortex—orchestrates interactions between the DMN and the EF system, dynamically transitioning between generative and evaluative modes ^61,63,64^. Supporting this gating function, multilayer network analyses demonstrate extensive reconfiguration among the EF system, DMN, and SN during creative idea generation, while evaluation engages a more integrated yet stable network architecture ^69^.

Alongside the modulation system, the EF networks —comprising primarily lateral prefrontal and posterior parietal cortices—provides top-down regulation to steer creative thinking toward task goals, such as response inhibition and idea revision that are critical for refining creative ideas. The EF system is particularly involved in suppressing dominant but less original responses, enabling individuals to consider novel, more creative alternatives ^68^. Such cognitive control engagement tends to vary with creative task demands ^70^, with flexible down-regulation often facilitating originality when dominant responses must be overcome ^44^. Creative thinking is characterized by heightened cooperation among these networks, with stronger coupling between the DMN and EF consistently predicting higher creative performance ^61,63^. Connectome-based predictive modeling reliably links enhanced functional connectivity across DMN, EF, and SN regions to individual differences in creative ability ^62,65,71^. Highly creative individuals exhibit greater network flexibility, showing more frequent transitions between the DMN and EF system ^64^ and increased temporal variability, especially within the DMN ^72^. How critical is the involvement of these cognitive control networks for the realization of creativity potentially impacted by interaction with ChatGPT, and even more differentiated in children and adults in the question of the current study.

### The current study

The current study aimed to determine the differences in cognitive engagement when children and adults are engaged while communicating with ChatGPT during a co-creative task. The study also examined whether the level of creativity was related to the level of cognitive control engagement. We hypothesized that children would show a lower engagement of cognitive control networks and that a connection between creativity and cognitive control functional connectivity would be observed in adults, but not in children.

## Methods

### Participants

Fifteen-6-7 year-old children, all first graders (mean age: 6.44 years, SD=4 months, 11 maels) and 16 adults (mean age: 38.12 years, SD=7.25 years, 4 maels) from an average-above-average socioeconomic status participated in the current study. For both age groups, the following inclusion criteria were applied: (1) intact vision and hearing, (2) no history of psychiatric or neurological delays (3) intact non-verbal and verbal IQ (>85), (4) born term (>37 weeks gestational age), (5) no contradiction to MR scanning: claustrophobia, any metal object in the body, including cardiac pacemaker, dental braces, and cochlear implants. Participants underwent a comprehensive behavioral assessment of their EF and creativity abilities, as well as an fMRI scan. Children and their parents signed informed assent and consent forms, respectively, and adults signed written consents before participating. Data was collected at the Technion Human MRI research center (TechMRC) at the Technion. The protocol was approved by the Ministry of Health Helsinki committee (for children) and by the institutions’ ethical committee at the Technion. Participants were compensated for their participation.

### Study procedure

Once children and adults were recruited and consented to participate in the study, they were invited to the Educational Neuroimaging Group (ENIG) facility at the Technion, where their cognitive and creative abilities were assessed. Then, upon completion, families were invited to the MRI facilities at the Technion, where ChatGPT was set up on the MRI console, and the child’s voice and all communication were recorded. All conversations with ChatGPT were recorded and analyzed for semantic complexity, a factor previously linked to creativity (following ^73^).

### Behavioral measures

To verify intact nonverbal IQ, children completed the Test of Nonverbal Intelligence (TONI ^74^) which includes a completion of missing parts in given matrices. To evaluate language abilities, the vocabulary test from the Peabody Picture Vocabulary Test (PPVT ^75^) was used. For adults, equivalent tests were administered: the Matrix task and the vocabulary task from the Wechsler Adult Intelligence Scale (WAIS) ^76^.

Creativity was assessed using the divergent association task (DAT; ^77^, https://www.datcreativity.com/task), conducted outside of the scanner. In this task, participants were required to retrieve 10 words that are as unrelated as possible, without any time constraints. The semantic relations between the words were calculated with a higher value representing higher creativity abilities (more divergent thinking).

### Neuroimaging measures

#### Talking to ChatGPT paradigm

The fMRI data were collected using a Prisma Magneton 3 Tesla scanner (Siemens Healthineers, Erlangen, Germany). A unique infrastructure set-up at the TechMRC allowed the data collection during the communication wth ChatGPT: an internet connection to the audio-visual system in the scanner, a real-life speech recording while fMRI data is collected using Optoacoustics system (Optoacoustics Ltd., Mazor, Israel), an in-house noise reduction system developed at the Technion allowing a clear sound transmission of the ChatGPT audio stimuli, and another in-house developed algorithm which allows a noise elimination from the audio recording allowing a direct transcript of the verbal output of the child. After desensitization and acclimation to the scanner environment (following our previous experience in this age group ^78^), anatomical data (T1) was collected for coregistration purposes. Upon connecting to ChatGPT, children were informed that they were about to speak to the “smartest computer in the world,” also known as ChatGPT, whereas adults were simply told that they would be speaking to ChatGPT (see prompts in the supplemental materials). Both groups were told that their goal is to open a store; It can be any store they would like, it can be anywhere, for anyone. They need to consult with ChatGPT regarding the type of store for 5 minutes while the child speaks to ChatGPT. Note, the instruction was given only when they were in the scanner, immediately before the start of the scan. An audio recording of the entire conversation was conducted via Audacity software (https://www.audacityteam.org/) and the transcript of the conversation was generated using ChatGPT.

#### Neuroimaging data

Each fMRI session lasted 5 minutes exactly and included a Time Repetition (TR) of 1 sec and the echo time (TE) was 28 milliseconds. The imaging setup included a field of view (FOV) of 20 × 20 × 13.6 cm, with a 100 × 100 matrix and slices of 2 mm thickness. High-resolution T1 images were collected for each participant to align the functional images accurately. The T1 scan parameters were a TR of 1750 milliseconds, a TE of 2.8 milliseconds, an inversion time of 900 milliseconds, and a flip angle of 9°, with a field of view (FOV) of 22.4 × 22.4 × 16 cm, a 224 × 224 × 160 matrix, and a slice thickness of 1 mm. All scans included a total of 300 volumes per session.

#### Neuroimaging data processing

The fMRI data were corrected for geometrical distortion and signal intensity variation due to static B0 field variations in T2* weighted fMRI data using corresponding B0 field map images acquired in every participant. Rigid boundary-based registration ^79^ to a high-resolution 3D, T1-weighted anatomical volume acquired within the same session, was performed. Nonlinear warping of this anatomical volume into standard space using a pediatric template was performed using ANTs ^80^ ^81^. Functional MRI data were subjected to the same preprocessing strategy outlined in ^82–88^. Using a model-based approach, we constructed a model for the functional connections among brain regions involved in the reading network ^89–95^. This included the export of functional connectivity values for the selected networks, as well as EF, attention and modulation networks (CO, FP, memory, DAN, VAN, DMN, and Salience ^96^) using the CONN toolbox implemented in SPM 12 (^97^; http://www.nitrc.org/projects/conn/). Functional connectivity between brain regions was quantified by the temporal correlation between their spontaneous BOLD signal fluctuations ^98^ based on Power’s atlas ^99^ containing 264 putative functional parcels, covering cognitive, sensory, and language/auditory functional networks (see Figure 1 for the networks spatial masks). To control for motion artifacts, functional connectivity analysis included a rigorous quality control (QC) process, which involved calculating the mean framewise displacement (FD) by summing the absolute values of the derivatives of the translational and rotational realignment estimates (after converting the rotational estimates to displacement at a 50mm radius). The QC vector was correlated across participants and conditions with the pairwise functional connectivity correlations between all parcels to ensure that motion artifacts do not drive the connectivity (see ^100^). None of the volumes reached FD > 0.5mm, and hence segments were not removed from the data (per the acceptable definition of data lasting fewer than five contiguous volumes were flagged and removed; ^100,101^). In addition, motion parameters were included as covariates in the first-level analyses. Due to significant difference in FD in children (X = .306, SD = .06) and adults (X = .226, SD = .05), t(30) = 3.65, *p* < .001), FD was controlled for in all analyses involving neuroimaging data.

**Figure 1.**
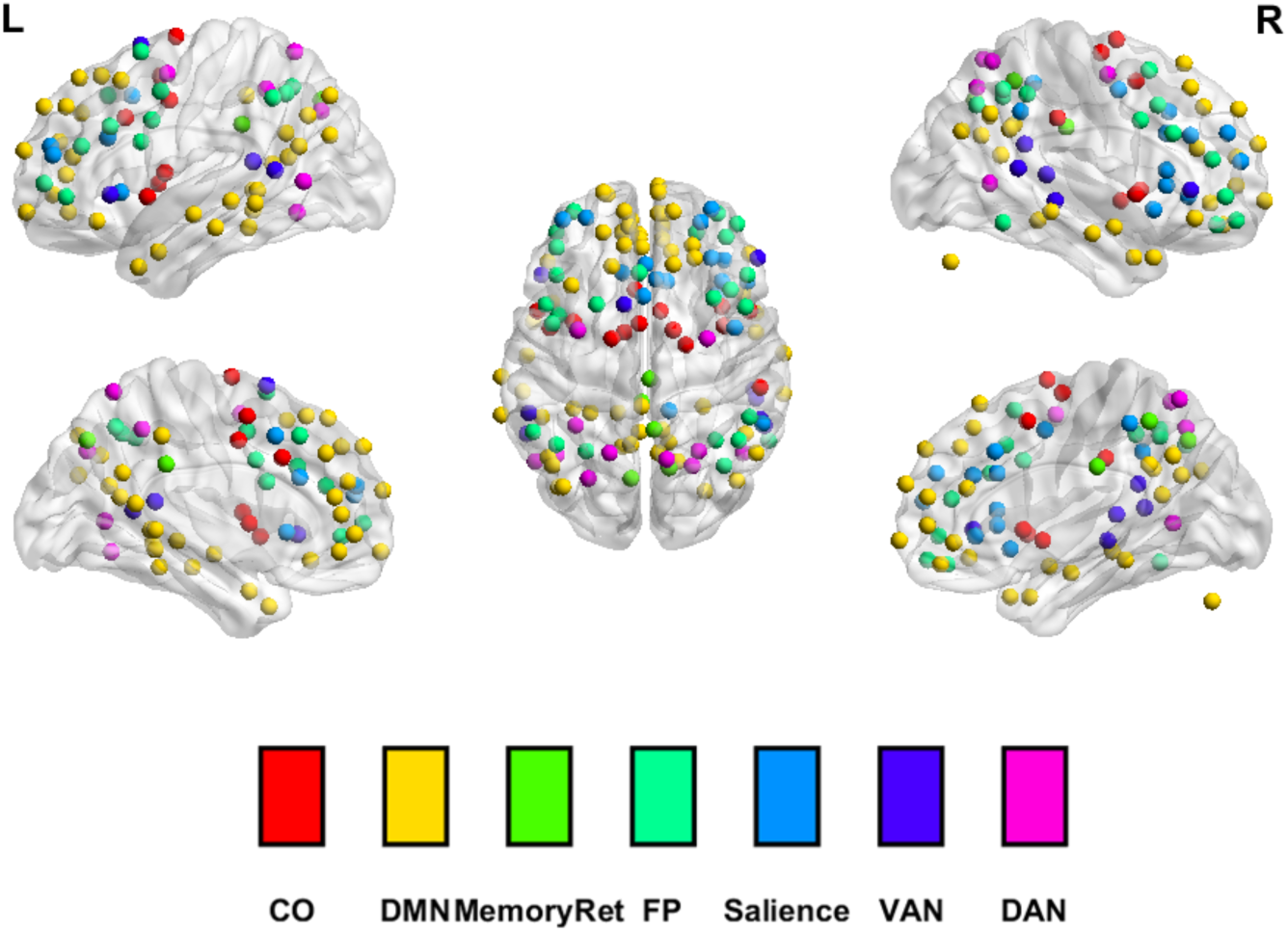
Executive functions, attention and modulation networks’ masks.

#### Divergent Sematic Integration (DSI) generated from the conversations with ChatGPT

DSI calculates the average semantic distance between all word pairs in a response using embeddings from a pre-trained model and cosine similarity. DSI has been shown to significantly predict originality on other creative tasks involving long-form text responses ^102,103^. Semantic-distance models work by placing words, sentences, or questions into a shared “semantic space,” where the distance between them reflects how similar or different their meanings are ^56^. For example, cat and dog will appear closer together in this space because they share many contextual similarities, while cat and democracy will be much farther apart. This is done in an unsupervised manner meaning they learn patterns of word co-occurrence and meaning directly from large text corpora without relying on predefined human ratings or labeled training data ^46^. DSI scores were computed for each participant for their entire discussion with ChatGPT based on their narrative transcripts. This was done to ensure enough text in each narrative to reliably compute DSI score. Thus, each DSI score represents the general diversity, or richness, of the conversation with ChatGPT. A higher score denotes a richer, more original narrative interaction with ChatGPT.

### Statistical analyses

#### Behavioral data analysis

Several *t*-test analyses were conducted to determine the differences in creativity levels assessed inside (DSI) and outside (DAT) of the scanner, among children and adults.

#### Neuroimaging data analysis

To determine differences in within- and between-network functional connectivity between children and adults, a multivariate analysis of variance (MANOVA) was conducted with group (children, adults) as a between-subjects factor and framewise displacement (FD) as a covariate to control for head motion. Dependent variables were mean Fisher-z-transformed connectivity values within seven canonical large-scale brain networks: cingulo-opercular (CO), default mode (DMN), memory retrieval, frontoparietal (FP), salience, ventral attention (VAN), and dorsal attention (DAN) networks, as well as their combinations. Univariate tests followed significant group effects and post hoc pairwise comparisons. Partial eta-squared (η²ₚ) values were reported as effect sizes.

#### Moderation analysis for the relationship between neuroimaging and behavioral data (creativity percentile scores)

To examine whether creativity moderates group differences in functional connectivity, we extended the above models by adding the creativity percentile score as a covariate and including the group-by-creativity interaction term.

Separate multivariate GLM models were estimated for both within- and between-network connectivity: 1) Model predictors: group (children vs. adults), creativity (centered), FD, and the group × creativity interaction, and 2) Dependent variables: within- or between-network connectivity measures as described above. For significant interactions, simple slope analyses and predicted values were examined to probe the direction and nature of the moderation effects.

## Results

### Behavioral *results*

Both children and adults demonstrated average verbal and nonverbal general abilities, with no significant differences between adults and children in these measures, as determined by independent-samples *t*-tests. Group comparisons revealed no significant difference in creativity performance (DAT) outside of the scanner between children and adults. However, creative interaction with ChatGPT (DSI) were significantly higher in adults compared to children (Table 1).

**Table 1.** Comparisons between children and adults in general abilities and creativity scores.

| Domain | Measure | Groups |  | t-test |
| --- | --- | --- | --- | --- |
|  |  | Children | Adults |  |
|  |  | Mean(SD) | Mean(SD) |  |
| Nonverbal abilities | Matrices | 10.41 (1.99) | 12.93(2.2) | -3.354, $p < .01$ |
| Verbal abilities | Vocabulary | 11.2(3.09) | 13.92(3.25) | -2.72, $p < .05$ |
| Creativity | DAT | 28.61 (19.59) | 26.16 (21.53) | 0.33, NS |
| | DSI (while speaking to ChatGPT) | .81 (.01) | .82 (.005) | -3.67, $p < .001$ |

Note. SS = Standard Score (M = 100, SD = 15), ScS = Scaled Score (M = 10, SD = 3) \*
$p < .05$ , \*\* $p < .01$ , \*\*\* $p < .001$ , n.s. = non significant

### Neuroimaging measures

#### Within networks analysis

After statistically controlling for head motion (FD), significant group differences in functional connectivity were observed in the FP network, *F*(1,27) = 8.77, *p* = .006, partial η² = .245, the salience network, *F*(1,27) = 5.65, *p* = .025, partial η² = .173, and the DAN, *F*(1,27) = 5.49, *p* = .027, partial η² = .169, with adults showing higher FC within these networks compared to children. These effects correspond to medium-to-large effect sizes, indicating that a substantial proportion of variance in connectivity was explained by group differences even after adjusting for motion.

**Table 2.** Within networks functional connectivity of EF, attention, and sensory networks.

| Network | Children<br>(Mean ± SD) | Adults<br>(Mean ± SD) | ANCOVA Results<br>($F(1,27)$, $p$, partial $\eta^2$) | Contrast |
| --- | --- | --- | --- | --- |
| Cingulo-Opercular | 0.307 ± 0.128 | 0.306 ± 0.106 | $F = 0.006$, $p = .940$,<br>partial $\eta^2 = .000$ | $C \approx A$ |
| Frontoparietal<br>(FP) | 0.113 ± 0.036 | 0.201 ± 0.085 | **$F = 8.773$, $p = .006$,**<br>**partial $\eta^2 = .245$** | **$A > C$** |
| Memory | 0.265 ± 0.168 | 0.342 ± 0.124 | $F = 1.938$, $p = .175$,<br>partial $\eta^2 = .067$ | $A > C$ |
| Default Mode<br>(DMN) | 0.181 ± 0.054 | 0.187 ± 0.057 | $F = 0.497$, $p = .487$,<br>partial $\eta^2 = .018$ | $A > C$ |
| Salience | 0.198 ± 0.073 | 0.289 ± 0.078 | **$F = 5.653$, $p = .025$,**<br>**partial $\eta^2 = .173$** | **$A > C$** |
| Dorsal Attention<br>(DAN) | 0.162 ± 0.098 | 0.211 ± 0.086 | **$F = 5.487$, $p = .027$,**<br>**partial $\eta^2 = .169$** | **$A > C$** |
| Ventral Attention<br>(VAN) | 0.369 ± 0.107 | 0.344 ± 0.120 | $F = 1.562$, $p = .222$,<br>partial $\eta^2 = .055$ | $C > A$ |

VAN: Ventral attention network; DAN: Dorsal attention network; FP: Fronto-parietal; CO:
Cingulo-opercular; DMN: default mode network; C: children; A: adults

#### Between Networks analysis

After statistically controlling for head motion (FD), several significant group differences emerged in between-network functional connectivity (Table 3). Compared with children, adults exhibited significantly stronger connectivity between the CO-FP networks, *F*(1,27) = 5.628, *p* = .025, η²ₚ = .173) and between the CO-Salience networks (*F*(1,27) = 8.182, *p* = .008, η²ₚ = .233. Similarly, adults showed greater coupling between the DMN-Memory, *F*(1,27) = 6.491, *p* = .017, η²ₚ = .194, and DMN-FP, *F*(1,27) = 5.570, *p* = .026, η²ₚ = .171. After correcting for multiple comparisons (using FDR correction ^104^, Benjamini–Hochberg, q = .05), the results for the FP findings remained significant.

**Table 3.** Between networks, functional connectivity of EF, attention, and sensory after controlling for FD.

| Network Pair | Children (Mean ± SD) | Adults (Mean ± SD) | ANCOVA Results ($F(1,27)$, $p$, $\eta^2_p$) | Contrast |
| --- | --- | --- | --- | --- |
| CO–DMN | 0.133 ± 0.070 | 0.119 ± 0.123 | $F = 0.006$, $p = .937$, $\eta^2_p = .000$ | C > A |
| CO–Memory | 0.015 ± 0.096 | 0.085 ± 0.110 | $F = 3.453$, $p = .074$, $\eta^2_p = .113$ | A > C |
| **CO–FP** | 0.043 ± 0.074 | 0.105 ± 0.052 | **$F = 5.628$, $p = .025$, $\eta^2_p = .173$** | **A > C** |
| **CO–Salience** | 0.109 ± 0.076 | 0.207 ± 0.079 | **$F = 8.182$, $p = .008$, $\eta^2_p = .233$** | **A > C** |
| CO–VAN | 0.165 ± 0.123 | 0.182 ± 0.103 | $F = 0.311$, $p = .582$, $\eta^2_p = .011$ | A > C |
| CO–DAN | 0.121 ± 0.077 | 0.116 ± 0.086 | $F = 0.494$, $p = .488$, $\eta^2_p = .018$ | C > A |
| **DMN–Memory** | 0.111 ± 0.045 | 0.150 ± 0.091 | $F = 6.491$, $p = .017$, $\eta^2_p = .194$ | **A > C** |
| **DMN – FP** | 0.090 ± 0.039 | 0.115 ± 0.065 | $F = 5.570$, $p = .026$, $\eta^2_p = .171$ | **A > C** |
| DMN–Salience | 0.111 ± 0.060 | 0.115 ± 0.081 | $F = 0.480$, $p = .494$, $\eta^2_p = .017$ | A > C |
| DMN–VAN | 0.196 ± 0.072 | 0.188 ± 0.091 | $F = 0.015$, $p = .902$, $\eta^2_p = .001$ | C > A |
| DMN–DAN | 0.116 ± 0.073 | 0.118 ± 0.086 | $F = 0.896$, $p = .352$, $\eta^2_p = .032$ | A > C |
| **Memory–FP** | 0.079 ± 0.064 | 0.104 ± 0.096 | $F = 6.449$, $p = .017$, $\eta^2_p = .193$ | **A > C** |
| Memory–Salience | 0.049 ± 0.059 | 0.079 ± 0.103 | $F = 3.990$, $p = .056$, $\eta^2_p = .129$ | A > C |
| **Memory–VAN** | 0.091 ± 0.073 | 0.133 ± 0.152 | $F = 5.514$, $p = .026$, $\eta^2_p = .170$ | **A > C** |
| **Memory–DAN** | 0.086 ± 0.076 | 0.144 ± 0.100 | $F = 5.944$, $p = .022$, $\eta^2_p = .180$ | **A > C** |
| FP–Salience | 0.138 ± 0.040 | 0.180 ± 0.060 | $F = 2.434$, $p = .130$, $\eta^2_p = .083$ | A > C |
| **FP–VAN** | 0.095 ± 0.059 | 0.167 ± 0.088 | $F = 5.820$, $p = .023$, $\eta^2_p = .177$ | **A > C** |
| **FP–DAN** | 0.039 ± 0.059 | 0.126 ± 0.043 | $F = 13.252$, $p < .001$, $\eta^2_p = .329$ | **A > C** |
| Salience–VAN | 0.012 ± 0.000 | 0.189 ± 0.007 | $F = 1.687$, $p = .205$, $\eta^2_p = .059$ | A > C |
| Salience–DAN | 0.010 ± 0.005 | 0.145 ± 0.005 | $F = 1.824$, $p = .188$, $\eta^2_p = .063$ | A > C |
| VAN–DAN | 0.002 ± 0.007 | 0.186 ± 0.007 | $F = 0.338$, $p = .566$, $\eta^2_p = .012$ | A > C |

VAN: Ventral attention network; DAN: Dorsal attention network; FP: Fronto-parietal; CO: Cingulo-opercular; DMN: default mode network; C: children; A: adults

Enhanced between networks connectivity in adults was also observed between the Memory Retrieval network and multiple systems, including the FP, *F*(1,27) = 6.449, *p* = .017, η²ₚ = .193, VAN, *F*(1,27) = 5.514, *p* = .026, η²ₚ = .170, and DAN *F*(1,27) = 5.944, *p* = .022, η²ₚ = .180. Finally, significant group effects were found for connectivity between the FP-VAN, *F*(1,27) = 5.820, *p* = .023, η²ₚ = .177, and FP-DAN, *F*(1,27) = 13.252, *p* = .001, η²ₚ = .329, with adults again showing higher connectivity.

Overall, these results suggest a broader and more integrated pattern of between-networks connectivity in adults relative to children, particularly involving the FP, DAN,VAN, memory, and salience systems. After correcting for multiple comparisons (FDR correction ^104^, Benjamini–Hochberg, q = .05), the results for the FP-salience and FP-DAN findings remained significant (Table 3). See also Figure 2.

**Figure 2.**
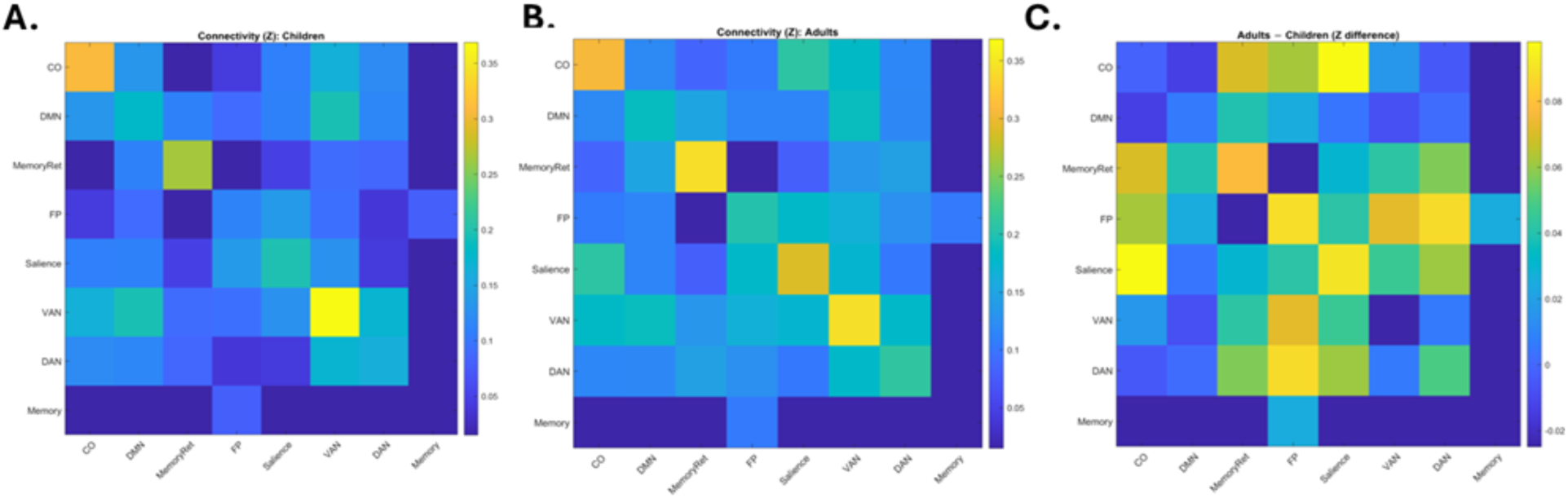
Averages for within and between networks’ functional connectivity (Z scores) The Z-score averages for within and between networks functional connectivity for children (A), adults (B) and the difference between the groups (C). The hot color represents a higher Z values whrease the cool color represents lower Z values.

### Moderation results

In order to determine whether creativity scores moderated the group effect on within-network connectivity (controlling for FD), we conducted moderation analyses, which revealed no significant group × creativity interaction effects across any network (p > .05).

Moderation analyses revealed that creativity outside the scanner (DAT) significantly interacted with group in predicting between-network functional connectivity. Specifically, significant moderation effects emerged for the FP–Salience, *F*(1,26) = 5.869, *p* = .023, partial η²ₚ = .184, and Salience–VAN, *F*(1,26) = 4.575, *p* = .042, partial η²ₚ = .150, pairs, indicating that the relationship between creativity and connectivity strength differed between children and adults for these networks.

To determine which group (children vs adults) shows a greater association between creativity skills (DAT) and functional connectivity between FP-Salience networks, an indepth slope analysis for the significant interactions was conducted, and revealed that adults’ show a positive significant slope (slope B = +0.028 per 1 SD creativity, *p* = .022, 95% CI [0.004, 0.052]) whrease children demonstrate a negative non significant slope (B = −0.014, *p* = .293, 95% CI [−0.042, 0.013] (n.s.)). Similar directions were found for Salience–VAN networks as well (Adults: B = +0.037, *p* = .062, 95% CI [−0.002, 0.077]; Children: B = −0.025, *p* = .273, 95% CI [−0.070, 0.021] (n.s.)). Overall, higher creativity is associated with stronger FP–Salience coupling in adults, but not in children (Figure 3).

**Figure 3.**
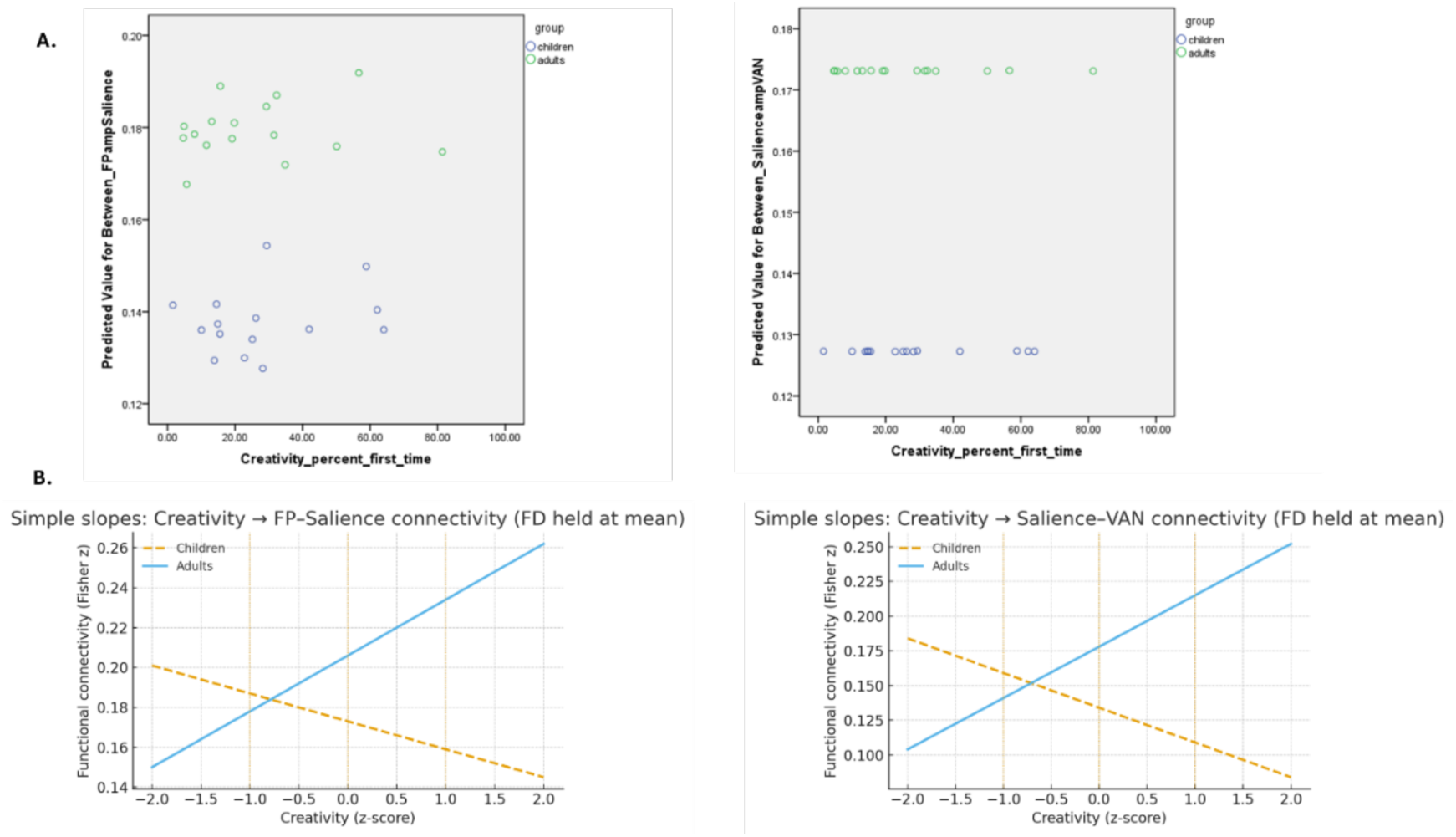
Association between creativity, functional connectivity and group differences. A. Correlation analysis between the finctional networks and creativity score. B. Slope differences analysis, comparing children to adults.

### Power analysis

Power calculations were anchored to published between-group differences in large-scale network connectivity. In younger versus older adults, Avelar-Pereira et al. reported a robust task-state difference in left FP-DAN coupling during the Multi-Source Interference Task, with groups of 29 and 30 and a large effect (Cohen’s d ≈ 1.3)^105^. In a separate high-versus low-creative adult comparison (12 per group), Beaty et al. reported ROI-to-ROI group differences spanning t values of 2.38–3.75 for IFG–DMN links, corresponding to large effects (d ≈ 1.0–1.5)^106^. To adopt a conservative planning assumption, we powered a priori for a moderate effect (Cohen’s d = 0.60–0.70), which requires approximately 44–33 participants per group for 80% power at α = .05 (two-sided). Our achieved sample (children, n = 15; adults, n = 16) is therefore well-powered for large effects, consistent with prior literature, but underpowered for moderate effects, guiding the interpretation of null or small effects accordingly.

## Discussion

How old do you need to be to use generative AI? With the invasion of these tools into our lives and even into the educational system ^107^, the current study aimed to explore the differences in cognitive control engagement when children vs. adults were orally communicating with ChatGPT during a creative task. In line with our hypothesis, children showed a lower engagement of cognitive control networks while communicating with ChatGPT compared to adults, despite an equal creativity level (age normalized) across groups. Moreover, and in line with our results, positive relations between creativity levels and cognitive control functional connectivity strength were observed in adults, but not in children.

### Reduced cognitive control, attention, and modulation networks during conversations with GenAI tools in children

Talking to ChatGPT or any other generative AI tools provides a unique opportunity to quickly gather information and build upon it to generate more higher-level, creative ideas. Whereas knowledge was previously gathered using search engines or other printed paper sources, today, gathering basic information is faster, thanks to the efficiency prompts provided by the user^108^. This process demands planning the utterance, monitoring your response and ChatGPT’s response, reorienting one’s attention to unpredictable responses, and maybe inhibiting responses when the conversation takes a different direction -these abilities all fall under the EF system definition ^109^. Whereas these stages may not dramatically differ in children and adults during human-to-human interaction, the speed of information flow and responses from ChatGPT and the lack of gestures usually received from humans but equipped with other verbal gestures^110^, might make this task very different than communicating with humans.

It was previously suggested that using generative AI tools results in lower alpha and beta waves generated from EEG data, with the claim that these tools demand less cognitive effort than an actual active search (engines vs self-search with no external tools) ^111^. The current study suggests that where adults do show an engagement of cognitive control (FP), attention (DAN), and modulation (salience) systems while speaking to ChatGPT, children show reduced functional connections within these networks. These findings are in line with developmental theories previously suggested ^112^.

Children also demonstrated reduced functional connectivity between the selected cognitive networks (FP-salience, FP-DAN, DMN-memory). In line with the reported increased between-networks connectivity during development ^34^, it appears that, when communicating with ChatGPT, children are less capable of retrieving information and internal mentation (DMN-memory), or adapting the behavior/thoughts in response to changes in a given stimulus (FP-DAN). These results, combined with studies demonstrating a lack of maturation of the cognitive control system at the age of 6 years ^35^ (first graders, the age of the participants in the current study), suggest that speaking to ChatGPT, which seems not to be so complicated, actually demands cognitive control, which is lower in children who participated in our study. This is especially true as the children in the current study are in their first grade, and there is a debate regarding the most appropriate age for using ChatGPT in school. Additional studies are needed to determine the exact time window in development when children gain the most from interacting with ChatGPT, also in terms of cognitive demands.

### Differential engagement of cognitive control networks in creative thinking: children versus adults

Creativity, measured in this study as DAT ^77^, is related to cognitive control, with evidence demonstrating increased creative abilities with increased frontal activation (specifically the Dorsolateral prefrontal cortex) from adolescence to young adulthood ^112^. Interestingly, although children and adults in the current study exhibited comparable levels of creativity measured outside of the scanner (age-normalized DAT), only adults showed significant associations between functional connectivity among cognitive control, attention, and modulation networks (FP–salience, salience–VAN) networks involved in the switching of executive functions and attention ^113^ critcal for creative thinking ^62–64,69,114^. In contrast, no such relations were observed in children.

Interestingly, previous research has linked creative thinking to the involvement of the salience network, which identifies and filters important or novel ideas before relaying them to the executive control system for goal-directed processing ^62,63,114^. Notably, the DMN, proposed as a key component in models of creativity ^66,115^ that support spontaneous thought and flexible memory retrieval, did not differ between adults and children, either overall or in relation to their DAT scores. This lack of difference may reflect the novelty of interacting with ChatGPT for both groups.

An additional possible explanation for these findings between-networks connectivity results and the absence of relations to creativity measures in children might be that children still do not have the neurobiological infrastructure to benefit from the conversation with ChatGPT, which also aligns with the lower DSI scores from their generated ChatGPT script. This is critical given the extended attention given to co-creative human-AI interaction ^14^, and their potential costs ^21–23,25^ and benefits ^17–20^ on human creativity. Future studies should examine the interaction between familiarity with ChatGPT and DMN involvement to determine whether the absence of engagement of DMN in relation to creativity is due to the equal novelty shared among children and adults using ChatGPT.

## Limitation

The results of the current study should be considered in light of the following limitations. First, the instructions for both age groups were slightly different: children were told they’d speak to “the smartest computer in the world,” whereas adults were simply told “ChatGPT.” In addition, the speech behavior, i.e., the conversation features such as turn count, words/min, response latency, were not accounted for and might also drive the results of the study (i.e., higher engagement and back and forth, might explain the greater cognitive control engagement). Moreover, there are apparent differences in vocabulary knowledge that change across the lifespan ^116,117^. Such a difference, which was not controlled here, might impact the co-creative interaction within the scanner and provide an alternative explanation for our results. Future studies are needed to replicate and extend our results, for example, by controlling for vocabulary knowledge in a similar study design. Finally, Post-hoc power analysis indicated 80% power to detect medium-to-large effects (Cohen’s d ≥ 0.73) for primary between-group comparisons. The moderation analyses were exploratory and likely underpowered for interaction effects, which typically require larger samples. Future studies with a sample size of ∼ 50 children per group are warranted to adequately test creativity × group interactions.

## Conclusions

This novel study examined the level of cognitive engagement of children while speaking to ChatGPT compared to adults, as well as how this cognitive engagement is related to creativity. The results point to the overengagement of the self-regulated networks in 6-year-old children, which warrants additional longitudinal studies to determine the turning point where cognitive control is more engaged and that creative thinking is related to cognitive control networks, similarly to adults. Given the rapid emergence of AI in our daily lives, alongside increased discussion on the costs and benefits of AI on creativity, our results highlight the significance of between-network functional coupling during human-AI creative acts. Decision makers, especially in the education system, as well as those related to children’s health (pediatricians), should pay particular attention to the official age when children can use ChatGPT, similarly to the attention given to screen exposure.

## Acknowledgement

The authors would like to thank the children, the parents, and the students participating in the current project. The authors would also like to thank the research team and students at the Educational Neueoimahging Group (of 2025) for helping with data collection and tasks’ administration. The authors would also like to thank Tuval Raz for his help in analyzing the transcripts of participants’ conversations with ChatGPT in the scanner.

## Author contribution

THK: Conceptualization, data collection, funding, analysis, wrote the first version of the paper, reviewed the final version of the paper

RF: Analysis, wrote the first version of the paper, reviewed the final version of the paper

YK: Analysis, wrote the first version of the paper, reviewed the final version of the paper

DL: Data collection, reviewed the final version of the paper

EA: Data collection, reviewed the final version of the paper

## Supplemental materials

### Prompt for children

You are having a conversation with a six-year-old Hebrew-speaking child.

The child will ask you questions related to opening a store, kindergarten, or school that they wish to start.

You should always reply in **only one sentence**, and **end your sentence with a question** to encourage the conversation.

The conversation should last **five minutes**.

Do **not** end the conversation on your own initiative.

Do **not** start until I tell you to begin — first, the child will say “hello / hi” for a test, and you should just reply “hello” in return.

Wait for my instruction to start.

### Prompt for adults

You are having a conversation with a Hebrew-speaking adult.

The adult will ask you questions related to opening a store, kindergarten, or school that they wish to start.

You should always reply in **only one sentence**, and **end your sentence with a question** to encourage the conversation.

The conversation should last **five minutes**.

Do **not** end the conversation on your own initiative.

Do **not** start until I tell you to begin — first, the adult will say “hello / hi” for a test, and you should just reply “hello” in return.

Wait for my instruction to start.

